# Interplay of Virulence Factors and Signaling Molecules: Albumin and Calcium-Mediated Biofilm Regulation in *Bordetella bronchiseptica*

**DOI:** 10.1101/2024.10.23.619887

**Authors:** Sabrina Laura Mugni, Nicolás Ambrosis, George A. O’Toole, Federico Sisti, Julieta Fernández

## Abstract

*Bordetella bronchiseptica*, a respiratory pathogen capable of infecting various mammals, including humans, is associated with chronic infections, contrasting with the acute infections caused by *Bordetella pertussis*. Both pathogens can form biofilm-like structures *in vivo*, providing tolerance against environmental stresses. Biofilm formation by *B. bronchiseptica* is regulated by the BvgAS two-component system, with intermediate concentrations of certain modulators inducing a phase favoring biofilm formation. Recent studies have highlighted the role of cyclic diguanylate monophosphate (c-di-GMP) in this process: elevated c-di-GMP levels stimulate biofilm formation, whereas phosphodiesterase (PDE) activation reduces biofilms. Respiratory secretions, which contain albumin and calcium at higher concentrations than standard growth media, promote an increase in the amount and extracellular localization of the adenylate cyclase toxin (ACT), an important *Bordetella spp*. virulence factor. Secreted ACT present in the extracellular media or attached to the outer membrane inhibits biofilm formation. Based on these observations, we hypothesized that serum albumin and calcium together inhibit biofilm formation and explored the potential role of c-di-GMP in this process. Our findings demonstrate that serum albumin and calcium inhibit *B. bronchiseptica* biofilm formation by two apparently independent mechanisms, increasing AC secretion and inducing c-di-GMP degradation. This study contributes to the understanding of the mechanisms governing *B. bronchiseptica* biofilm formation and its modulation by host factors.

## INTRODUCTION

*Bordetella bronchiseptica* is a respiratory pathogen known to infect a variety of mammals, including humans. Infections caused by this pathogen are characterized by chronic and persistent infections, contrasting with the more acute infections caused by the human-exclusive pathogen *B. pertussis*. It is increasingly accepted that both pathogens can survive asymptomatically in the upper respiratory tract for extended periods (1–3). This asymptomatic persistence may explain the difficulty in eradicating *B. pertussis* despite high vaccine coverage.

Biofilm formation by *B. bronchiseptica* is a highly regulated process controlled in part by the two-component system BvgAS, which also regulates the expression of virulence factors (4). While the BvgAS system is believed to be constitutively active, its activity can be modulated in the laboratory by millimolar concentrations of nicotinic acid or magnesium sulfate. Intermediate concentrations of these modulators can induce an “intermediate phase” in *B. bronchiseptica* where several virulence factors are absent, yet biofilm formation is maximized. Recent investigations by our group have highlighted the role of the second messenger cyclic diguanylate monophosphate (c-di-GMP) in regulating biofilm formation in *B. bronchiseptica*, particularly during the intermediate phase (5).

Elevated levels of c-di-GMP, resulting from the activation of diguanylate cyclases (DGC), stimulate biofilm formation, whereas activation of phosphodiesterases (PDEs) leads to c-di-GMP degradation and subsequent biofilm reduction. The BrtA/Lap system, responsive to c-di-GMP levels, is required for biofilm formation in the intermediate phase (6). BrtA, a large adhesin, remains attached to the outer membrane until its cognate protease, LapG, cleaves the N-terminal periplasmic domain of this adhesin. The release of BrtA is associated with low biofilm levels. LapG, a periplasmic protease, is sequestered by LapD when cytosolic c-di-GMP concentrations are high, preventing BrtA cleavage. Hence, if c-di-GMP is elevated, BrtA remains on the bacterial surface and biofilm is formed and stabilized.

Both *B. pertussis* and *B. bronchiseptica* can form biofilm-like structures *in vivo* on the respiratory epithelium (7, 8). Biofilm formation is typically described as a tolerance mechanism versus antibiotics and other environmental stresses. The biofilm matrix of *B. bronchiseptica* is comprised of proteins, polysaccharides, and extracellular DNA (eDNA), which provide structural integrity and protection for the bacterial community (9). Key adhesins such as filamentous hemagglutinin (FHA) play pivotal roles in biofilm formation and can be secreted into the extracellular milieu (10). Interestingly, the presence of ACT has been found to diminish biofilm formation (11). Deleting the *cyaA* gene encoding the ACT of *B. bronchiseptica* increases biofilm formation, while the addition of the ACT to the extracellular medium inhibits biofilm formation in *Bordetella pertussis* (4, 11).

Respiratory secretions, which contain albumin and calcium, promote an increase in the amount and membrane localization of active ACT (12). This increased secretion and surface localization of ACT can be elicited *in vitro* by both serum and albumin. The response to albumin is not mediated through the regulation of ACT at the transcriptional level or activation of the BvgAS two-component system (12). Considering the promotion of ACT secretion by serum albumin and its reported involvement in biofilm formation, we hypothesized that serum albumin may inhibit biofilm formation. Furthermore, given the regulatory role of c-di-GMP in biofilm formation, we aimed to investigate whether this second messenger is involved in the signaling triggered by serum albumin.

Here we demonstrate that bovine serum albumin (BSA) and calcium inhibit the formation of *B. bronchiseptica* biofilms. While the FHA adhesin does not appear to be involved in this process, ACT is implicated, with more ACT associated with reduced biofilms. Additionally, we show that the surface localization of the BrtA adhesin, responsible for biofilm formation in the intermediate phase, is regulated by BSA and calcium through the activation of PDEs, which in turn reduce c-di-GMP levels. Through this study, we seek to deepen our understanding of the mechanisms underlying *B. bronchiseptica* biofilm formation and its modulation by host factors.

## RESULTS

### Albumin and Calcium Inhibit B. bronchiseptica Biofilm Formation

Gonyar and colleagues demonstrated that albumin and calcium present in host serum were responsible for an increase in ACT secretion into the extracellular media (12). Additionally, ACT present in the extracellular media, and possibly on the external surface of the bacteria, inhibits biofilm formation (11). Taking these two results together, we hypothesized that albumin and calcium at serum concentrations would inhibit biofilm formation. To test this hypothesis, we evaluated biofilm formation using the crystal violet (CV) technique in the absence and presence of albumin and calcium. Following the methods of Gonyar *et al*., we used bovine serum albumin (BSA) to test the effect of albumin, as their experiments showed no differences between the two additions regarding an impact of biofilm formation.

As shown in Figure 1, a significant inhibition of biofilm formation was noted in the presence of BSA+Ca at serum-relevant concentrations for *B. bronchiseptica* 9.73. Reduction was not attributable to differences in growth (Fig. S1). The effect is also observed in *B. bronchiseptica* strain RB50 (Fig. 1A). Inhibition was not observed when BSA was pretreated with proteinase K, thus ruling out potential effects of contaminants present in the BSA (Fig. 1B).

**Figure 1.**
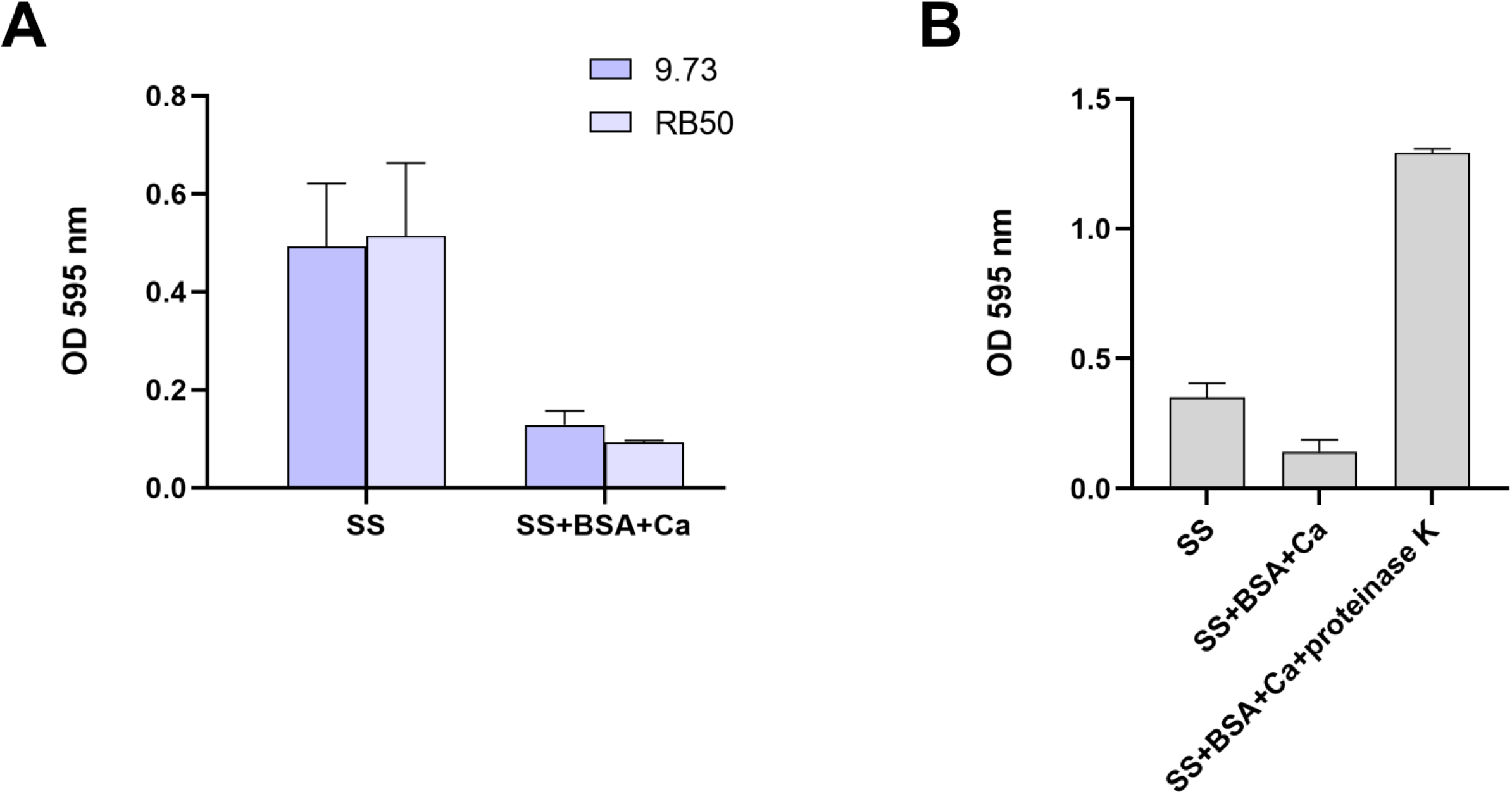
Albumin and Calcium Inhibit *B. bronchiseptica* biofilm formation. Biofilm formation on PVC 96-well plates was assessed of overnight cultures of wild type *B. bronchiseptica* 9.73 (A) or RB50 (B) grown in SS or SS supplemented with 2.0 mg/ml BSA and 2.0 mM CaCl_2_. C. Biofilm formation of *B. bronchiseptica* in SS supplemented with BSA+Ca pretreated with proteinase K. Visualization and quantification were performed using the crystal violet (CV) staining technique. Results are the average of at least three independent experiments. ** indicate significant differences p<0.01 (ANOVA).

*B. bronchiseptica* biofilm formation is regulated by the two-component system BvgAS. When partially active, meaning in the intermediate phase, *B. bronchiseptica* forms the maximum biofilm. To determine if BSA+Ca inhibition is also observed in other phases besides the virulent phase, we conducted static biofilm assay experiments with bacteria modulated in intermediate and avirulent phases using nicotinic acid (NA). As previously described, biofilm formation was higher in intermediate phase, that is, in the presence of 0.5 to 2 mM NA. In all NA concentrations tested, BSA+Ca inhibited biofilm formation (Fig. 2A).

**Figure 2.**
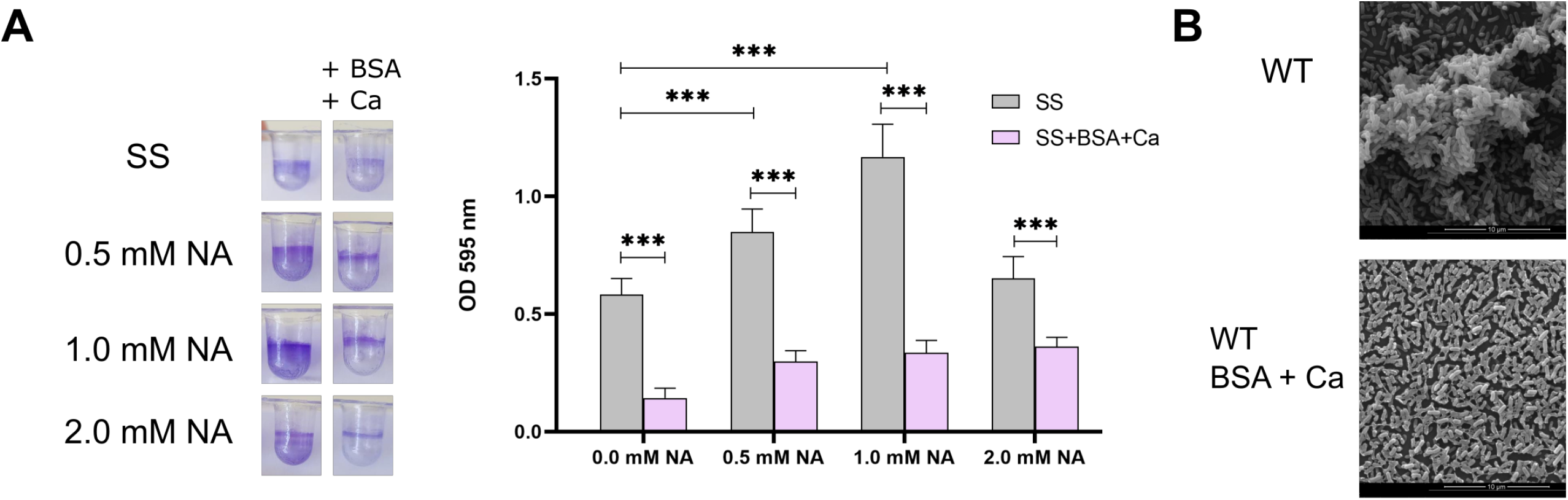
The Effect of BSA+Ca on Biofilm Formation Depends on the Virulent Phase. A. Wild type *B. bronchiseptica* was grown for 24 hours in SS or SS supplemented with 2.0 mg/ml BSA and 2.0 mM CaCl_2_ with the indicated final concentration of added nicotinic acid (NA). Biofilm was stained with CV and quantified after resuspension in 33% v/v acetic acid. Results are the average of at least three independent experiments. *** indicates significant differences versus SS at the same nicotinic acid concentration (p<0.001. ANOVA). B. Scanning electronic microscopy of 24-h cultures of wild type *B. bronchiseptica* grown in SS with 2.0 mM NA or in SS with 2.0 mM NA, 2.0 mg/ml BSA and 2.0 mM CaCl_2_.

Biofilm inhibition in the intermediate phase was confirmed by scanning electron microscopy (SEM) (Fig. 2B). We chose the intermediate phase for SEM observation because it was the condition where biofilm persisted even in the presence of BSA+Ca. When *B. bronchiseptica* was grown in SS with 2 mM NA, it formed three-dimensional structures, but in the presence of BSA+Ca, only a monolayer was observed (Fig. 2B).

In conclusion, our results confirm our initial hypothesis that host serum components, albumin and calcium, inhibit *B. bronchiseptica* biofilm formation.

### Filamentous Hemagglutinin is not Required for the Reduction in Biofilm Formation by Added BSA and Calcium

FHA is an adhesin located at the bacterial surface, crucial for biofilm formation. FHA can also be cleaved and released into extracellular media by a tightly regulated process involving various proteases, including DegP (13, 14). Serra and co-workers reported inhibition of *B. pertussis* biofilm formation by exogenous addition of FHA (10), suggesting that interfering with FHA-mediated interactions, possibly through the production of proteolyzed FHA, could reduce biofilm formation. However, the impact of FHA DegP-mediated proteolysis and release from the cell surface on biofilm formation has not been evaluated. If the presence of FHA on the bacterial surface is necessary to establish bacterium-bacterium interactions leading to biofilm formation, BSA+Ca might disrupt this process by stimulating FHA release. If our hypothesis is correct, the deletion of *degP*, which abolishes FHA release, would reverse the negative effect of BSA+Ca on biofilm formation.

*B. bronchiseptica* with clean deletion in *degP* was evaluated for biofilm formation by CV. In the absence of the *degP* gene, BSA+Ca continued to reduce biofilm formation (Fig. 3A). Given that the effect of FHA on biofilm formation is most evident in SS medium with 0.8 mM NA, we next performed biofilm experiments under these conditions (SS + 0.8 mM NA). Deletion of the *degP* significantly impairs biofilm formation both in the absence and presence of BSA+Ca, suggesting that FHA released from the cell surface likely promotes biofilm formation (Fig. 3A). Differences were not due to a growth effect (Fig. S2). Dot blot analysis of intact cells using an anti-FHA antibody revealed a stronger signal in the BbΔ*degP* strain compared to the isogenic parent strain (Fig. 3B), indicating that the accumulation of FHA on the cell surface is linked to a reduction in biofilm formation. In conclusion, we found that biofilm inhibition by BSA+Ca occurs independently of FHA processing in *B. bronchiseptica*.

**Figure 3.**
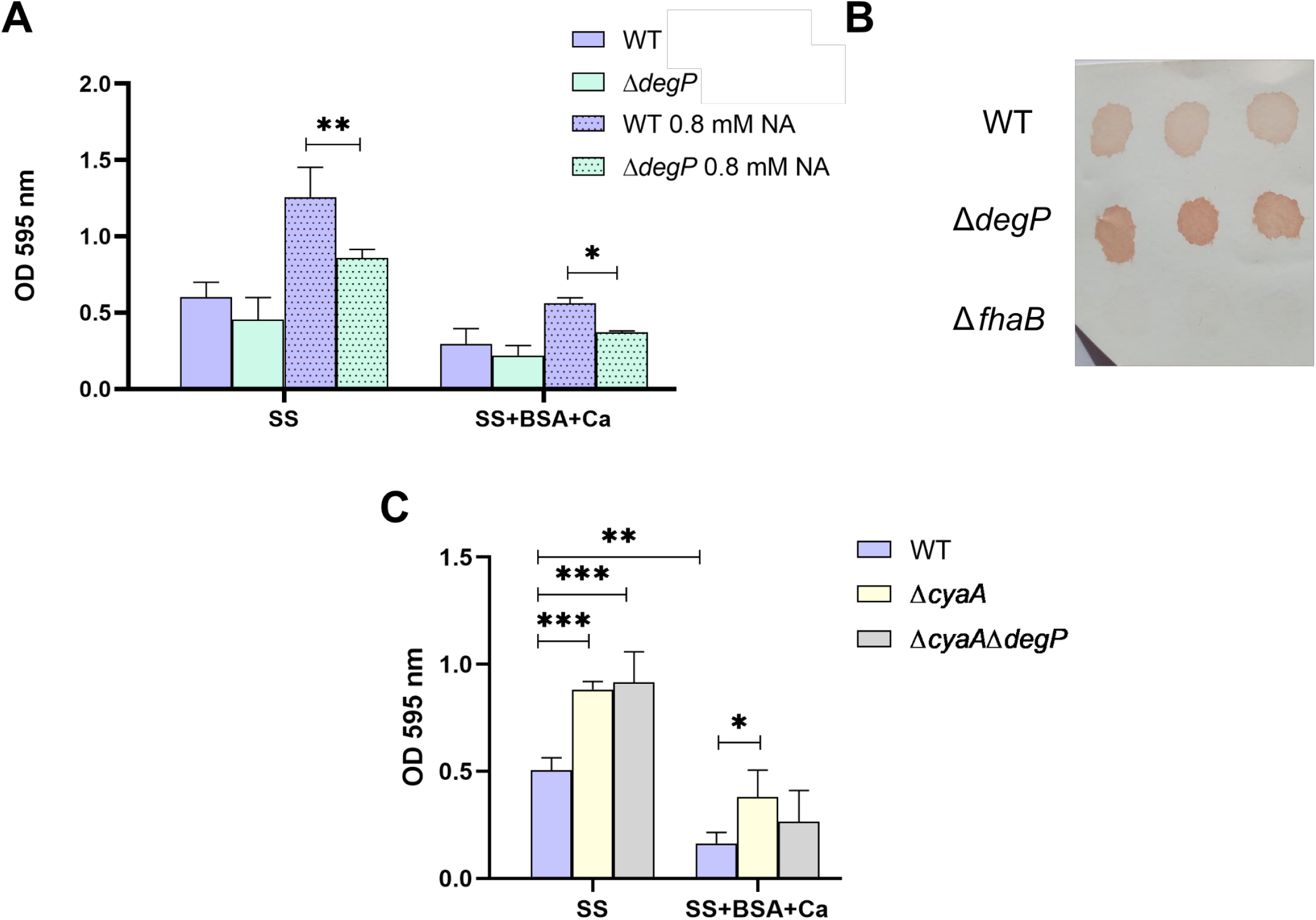
The Role of Filamentous Hemagglutinin and Adenylate Cyclase in Biofilm Formation. Biofilm formation on PVC 96-well of overnight cultures of wild type *B. bronchiseptica*, and *Bb*Δ*degP* (A) or *Bb*Δ*cyaA* and *Bb*Δ*cyaA*Δ*degP* (C) grown is SS or SS supplemented with 2.0 mg/ml BSA and 2.0 mM CaCl_2_. Biofilm was stained with CV and quantified after resuspension in 33% v/v acetic acid. Results are the average of at least three independent experiments. ** or *** indicate significant differences (p<0.01 and p<0.001 respectively. ANOVA). B. Cell surface levels of FHA as measured by dot blot. A representative dot blot assay with triplicates is shown.

### Adenylate Cyclase Toxin-Mediated Biofilm Inhibition by BSA+Ca

Adenylate cyclase toxin, via its AC domain, binds FHA on the cell surface thus preventing FHA interactions necessary for biofilm formation (11). Irie and co-workers reported increased biofilm formation in the absence of ACT (4). As described previously, albumin and calcium stimulate ACT secretion. Considering all these data, we hypothesized that the observed biofilm inhibition in the presence of BSA+Ca may be caused by increased secreted ACT. To test this idea, we first evaluated biofilm formation by a *cyaA* mutant (*Bb*Δ*cyaA*) to confirm the results obtained by Irie and co-workers (Fig. 3C). The *Bb*Δ*cyaA* mutant formed more biofilm than the wild-type strain in the virulent phase and also formed more biofilm than the WT in the presence of BSA+Ca (Fig. 3C). To rule out the possibility that FHA plays a role in biofilm reduction but that this effect was masked by the ACT effect, we evaluated biofilm formation in a double mutant strain lacking both *degP* and *cyaA*. No significant differences were observed between *Bb*Δ*cyaA*Δ*degP* and *Bb*Δ*cyaA* (Fig. 3C). Given that the strain lacking ACT showed less inhibition than the WT, these data indicate that ACT may be involved in biofilm inhibition by added BSA+Ca.

### BSA + Ca-Induced Decrease in c-di-GMP Levels Correlates with Biofilm Reduction

We previously described that c-di-GMP regulates biofilm formation in *B. bronchiseptica* (5). If biofilm inhibition were entirely or partially mediated by changes in c-di-GMP level, we predict that intracellular levels of this second messenger should decrease in the presence of BSA+Ca. To test this hypothesis, we evaluated c-di-GMP intracellular concentration in SS media and in SS supplemented with BSA+Ca. Considering that intracellular levels of c-di-GMP might be low in unstimulated conditions and a decrease may be difficult to detect, we also evaluated c-di-GMP levels in a strain overexpressing an active DGC (BdcA). We previously showed that this strain produces higher levels of c-di-GMP (15). As shown in Figure 4A, BSA+Ca addition resulted in a significant decrease in c-di-GMP intracellular levels of the WT and when the DGC BdcA is expressed from a plasmid, consistent with the reduced biofilm phenotype upon addition of BSA+Ca.

**Figure 4.**
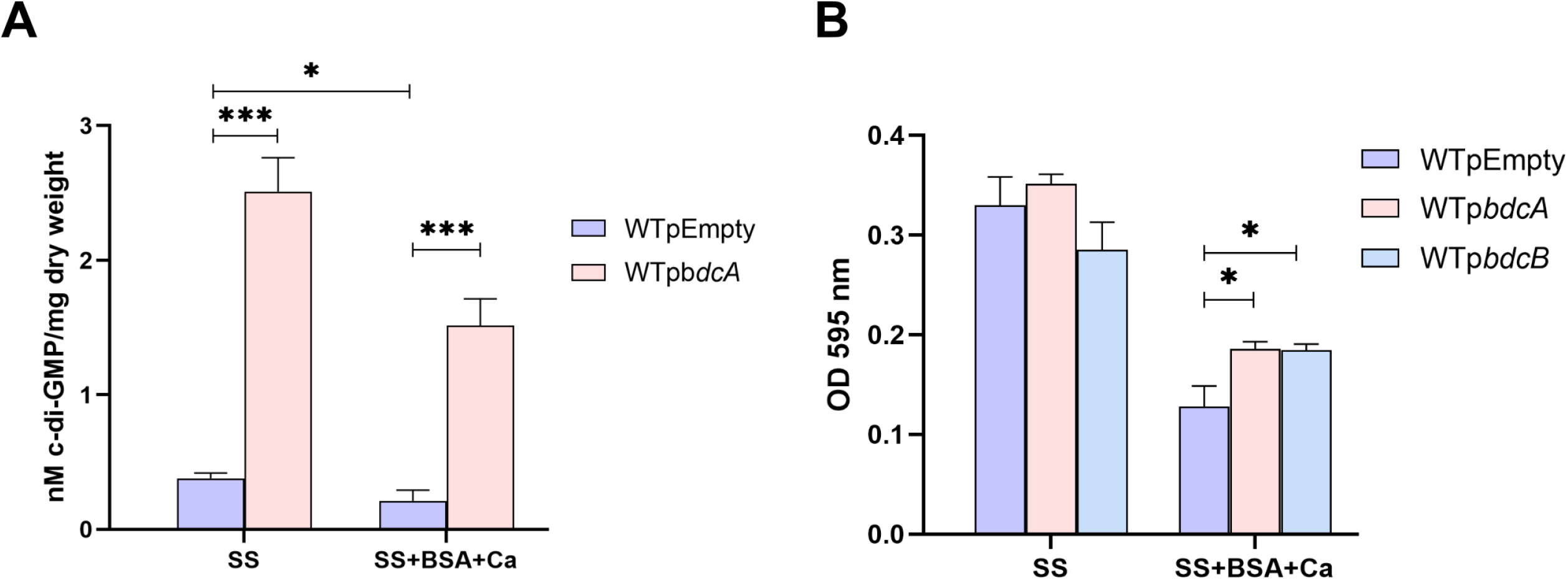
BSA+Ca-Induced Decrease in c-di-GMP Levels Correlates with Biofilm Inhibition. A. Intracellular c-di-GMP levels were measured in wild type *B. bronchiseptica* or *Bb*p*bdcA* cells grown in SS or SS supplemented with 2.0 mg/ml BSA and 2.0 mM CaCl_2_. Results are the average of three independent experiments. * or *** indicate significant differences (p<0.05 and p<0.01 respectively. ANOVA). B. Biofilm formation on PVC 96-well plates of overnight cultures of wild type *B. bronchiseptica, Bb*p*bdcA* and *Bb*p*bdcB* grown in SS or SS supplemented with 2.0 mg/ml BSA and 2.0 mM CaCl_2_. Biofilm was stained with CV and quantified after resuspension in 33% v/v acetic acid. Results are the average of at least three independent experiments. * indicate significant differences (p<0.05. ANOVA).

To test if these c-di-GMP levels correlate with biofilm formation, we evaluated biofilm formation of strains expressing DGCs from a plasmid. We expressed two different DGCs, BdcA and BdcB, which are membrane- and cytosolically-localized DGCs, respectively (5, 16). While BSA+Ca inhibited biofilm formation on both surfaces, biofilm was observed in both strains expressing a DGC from a plasmid (*Bb*p*bdcA* and *Bb*p*bdcB*) (Fig. 4B). This finding is consistent with our hypothesis that reduced c-di-GMP is associated with the reduction of biofilm formation by added BSA+Ca.

### Exploring PDEs and BvgR Influence on Biofilm Formation

As shown above, c-di-GMP levels are diminished in the presence of BSA+Ca, thus it is possible that one or more PDEs are activated upon addition of these compounds. Therefore, if these PDEs are deleted from the *B. bronchiseptica* genome, biofilm inhibition should not occur upon the addition of BSA+Ca.

*B. bronchiseptica* has nine putative proteins with a PDE-associated EAL domain encoded in its genome. Among these, LapD, BB2109, and BvgR have degenerate EAL domains, suggesting they likely lack PDE activity. BB2957 and BB3317 are dual-function proteins that may exhibit both DGC and PDE activities. Therefore, these proteins were excluded from our analysis. Among the remaining candidates, BB2110 (we propose the name <u>P</u>hospho<u>d</u>iesterase <u>B</u>, PdeB) is a bona fide EAL-containing protein. Multiple attempts to delete *bb2110* were unsuccessful. This narrowed our focus to three potential candidates: the two membrane-associated PDEs, PdeC and PdeD, and a cytosolic PDE, PdeA, previously described by our team (17). Although BvgR does not possess essential amino acids for PDE activity, we have described its importance in biofilm formation(18). Therefore, we included this protein in the analysis.

To test our hypothesis, we constructed individual mutations in each of these PDEs. All strains were evaluated for biofilm formation on a PVC surface. As previously described, the *Bb*Δ*bvgR* mutant strain formed more biofilm than the wild type in standard SS (Fig. 5A). Although the biofilm levels of the PdeC and PdeD mutants in SS appeared qualitatively lower than those of the wild type, no significant differences were observed. When grown in the presence of BSA+Ca, none of the PDEs mutants showed significant differences compared to the wild-type strain (Fig. 5A). However, a modest but significant increase in biofilm formation was observed in the *Bb*Δ*bvgR* mutant in the presence of BSA+Ca (Fig. 5A). We also deleted all four genes coding for PDEs in case of potential redundancy. The quadruple *Bb*Δ4PDE mutant showed no difference from the WT in standard SS medium, but the *Bb*Δ4PDE mutant showed significantly more biofilm formation than the wild-type strain in the presence of BSA+Ca (Fig. 5B), consistent with our hypothesis above. Taken together, BSA+Ca likely stimulates the production and/or activity of two or more PDEs to reduce cellular levels of c-di-GMP.

**Figure 5.**
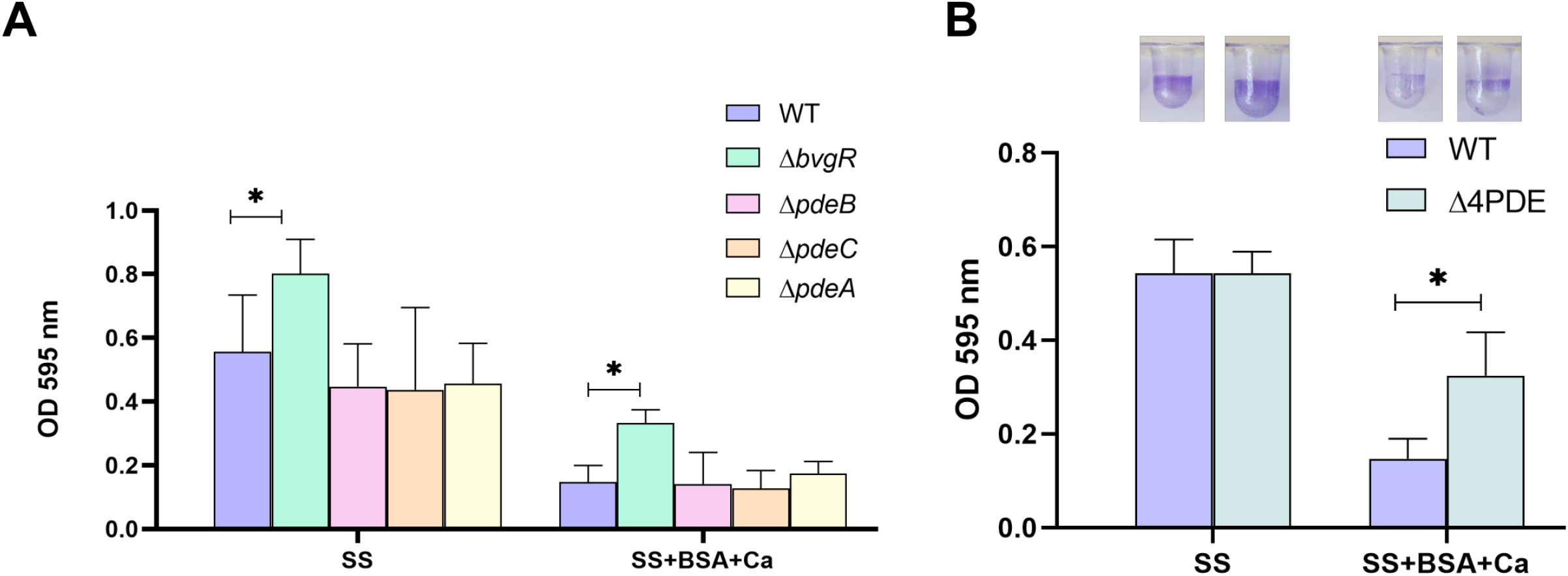
Exploring the Influence of PDEs and BvgR on Biofilm Formation. Biofilm formation on PVC 96-well plates of overnight cultures of wild type *B. bronchiseptica* and individual PDE mutants (A) or the *Bb*Δ4PDE multiple mutant (B) grown in SS or SS supplemented with 2.0 mg/ml BSA and 2.0 mM CaCl_2_. Biofilm was stained with CV and quantified after resuspension in 33% v/v acetic acid. Results are the average of at least three independent experiments. * or ** indicate significant differences ((p<0.05 and p<0.01 respectively. ANOVA).

### BSA+Ca Effect on BrtA Surface Levels and Biofilm Formation

The effect of BSA+Ca was observed in all virulent phases, but *B. bronchiseptica* biofilm formation peaks in the intermediate phase. Therefore, we decided to evaluate this phase for impacts on BrtA secretion and surface levels. Previously, we described BrtA as the adhesin responsible for biofilm formation in the intermediate phase (2.0 mM NA), and that the elaboration of this protein on the cell surface is regulated by c-di-GMP (6). Using a 2.0 mM concentration of NA, we assessed the effect of BSA+Ca on BrtA secretion. The supernatant of wild type bacteria in the presence of BSA+Ca exhibited significantly higher levels of BrtA, but no difference in whole cell levels (Fig. 6A). Moreover, the presence of BrtA on the bacterial surface, evidenced by a whole cell dot blot, was consistent with the increased secreted BrtA phenotype (Fig. 6A). The elevated BrtA in the supernatant and absence of this protein on the cell surface when cells were treated with BSA+Ca may explain the reduction in biofilm formation in the intermediate phase. If this hypothesis holds true, the inability to secrete BrtA would inhibit the effect of BSA+Ca. Deletion of the BrtA-specific protease LapG impedes BrtA release into the extracellular media (6). We evaluated biofilm formation of wild type and *Bb*Δ*lapG* on PVC and glass surfaces. As expected, based on our hypothesis, the LapG mutant did not exhibit a reduced capacity for biofilm formation in the presence of BSA+Ca. Moreover, biofilm formation, measured by CV, was observed to be greater than that of the WT strain. To better understand this phenomenon, we examined biofilm formation using SEM. As shown in figure 6C the LapG mutant in the presence of BSA+Ca formed three-dimensional structures.

**Figure 6.**
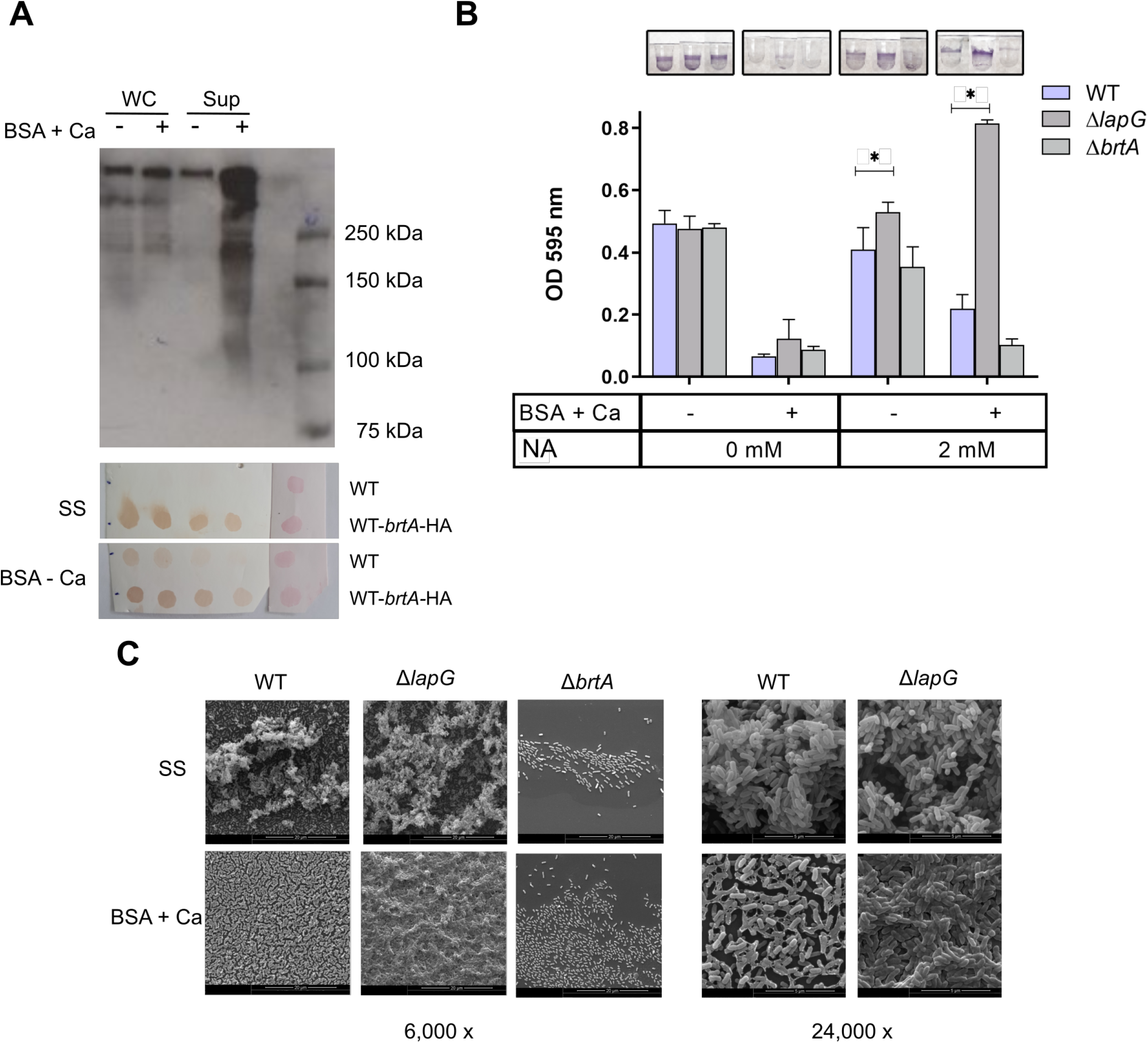
Effect of BSA+Ca on BrtA Secretion and Biofilm Formation. A. Analysis to determine BrtA presence in whole-cell (WC) lysates, supernatants or on the bacterial surface from late-phase growth culture of wild type *B. bronchiseptica* grown in SS or SS supplemented with 2.0 mg/ml BSA and 2.0 mM CaCl_2_. B. Biofilm formation on PVC 96-well plates of overnight cultures of wild type *B. bronchiseptica, Bb*Δ*lapG* and *BbΔbrtA* grown in SS or SS supplemented with 2.0 mg/ml BSA and 2.0 mM CaCl_2_ with the indicated concentrations of NA. Biofilm was stained with CV and quantified after resuspension in 33% v/v acetic acid. Results are the average of at least three independent experiments. * indicate significant differences (p<0.05. ANOVA). C. Scanning electronic microscopy of 24-h cultures of *B. bronchiseptica* presented in panel B in the presence of 2.0 mM NA.

## DISCUSSION

The infection process of *B. bronchiseptica* can be summarized as follows: initially, the bacteria colonize the respiratory epithelium, where they may persist asymptomatically for an extended period, facilitated by the formation of biofilm-like structures. The bacteria express important adhesins, including FHA and BrtA, and secrete eDNA, contributing to biofilm persistence. The local concentration of albumin and calcium in this zone may not be sufficient to trigger a biofilm disruption response. Under certain unknown conditions, the bacteria detach from the biofilm and progress towards the lower respiratory tract to the lungs. In this environment, they encounter concentrations of albumin and calcium sufficient to stimulate the secretion of ACT, resulting in biofilm inhibition. Additionally, in this situation, c-di-GMP levels would be low with loss of cell-surface BrtA, consistent with reduced biofilm formation. We have previously reported that high levels of c-di-GMP inhibit Type III secretion system (TTSS) activity. Therefore, in the presence of albumin and calcium, *B. bronchiseptica* in the lung would also be expressing the TTSS system, necessary to counteract the host’s immune response.

The results presented here support the proposed model above. The presence of albumin (mimicked by BSA in our experiments) and calcium induced an increase in extracellular ACT level and loss of cell-surface BrtA. The inability to produce extracellular ACT or release cell-surface BrtA partially prevented the inhibitory effect of BSA+Ca, demonstrating that this inhibition is mediated by multiple factors, including ACT, BrtA, and potentially other unidentified components. Although FHA is an important adhesin for biofilm formation, likely facilitating FHA-FHA bridges between bacteria, our experiments did not show that its proteolysis affects the action of BSA+Ca. However, when its release was blocked through the deletion of the *degP* gene, we observed that *B. bronchiseptica* could not properly form biofilm. This finding is particularly intriguing, as other studies have shown that exogenous FHA inhibits biofilm formation in the closely related *B. pertussis* (10).

Our findings also confirm the involvement of c-di-GMP in biofilm regulation by albumin and calcium. However, we were unable to establish a direct link between ACT levels and c-di-GMP, suggesting that albumin and calcium may independently impact ACT and BrtA. Previously, we reported that *B. bronchiseptica* overexpressing the DGC BdcA exhibited lower total ACT levels, indicating that a possible connection between ACT and BrtA secretion cannot be ruled out.

The second messenger c-di-GMP regulates various phenotypes through the activation of distinct PDEs and DGCs, each responsive to different signals, as recently proposed in the Bowtie model (19). Phenotype regulation is governed by global c-di-GMP concentrations, which result from the balanced activities of DGCs and PDEs. Henggé Fountain model proposes the existence of a general regulatory PDE that maintains low levels of c-di-GMP (20). According to this model, the removal of the master regulatory PDE would affect c-di-GMP levels and the responses to albumin and calcium, although it does not imply this PDE is specific to any particular phenotype. Another non-exclusive possibility is that the regulation of biofilm formation by c-di-GMP in response to albumin and calcium is local, as proposed in the Hub model (19). In this scenario, the phenotype would only be observed when the specific PDE is removed. In our study, we found that only BvgR may play a role in biofilm inhibition mediated by albumin and calcium. Interestingly, there is no evidence to support that BvgR is an active PDE (18). Further studies are needed to identify other factors involved in this regulation and to understand how the bacteria sense the presence of albumin and calcium.

There is evidence that high concentrations of albumin and calcium resemble the environment in the host’s nostrils. Infiltration of immune cells from nasal tissue to the site of infection transports significant amounts of albumin and calcium, which are present in the extracellular fluid. *Bordetella spp*., established as microcolonies on the epithelial surface, may sense this as a signal to disassemble biofilm or prevent the formation of new biofilm structures, facilitating its spread to other regions where the immune response has not yet developed.

Finally, we would like to emphasize an important consideration regarding the evidence that albumin and calcium inhibit biofilm formation *in vitro*. This inhibition occurs under conditions with high concentrations of albumin and calcium, whereas typical media used in *in vitro* experiments contain significantly lower levels of both, perhaps explaining why these phenotypes have not been observed previously.

## MATERIAL AND METHODS

### Bacterial strains and growth conditions

*B. bronchiseptica* strains (Table S1) were grown at 37°C either on Bordet-Gengou agar (BGA, Difco) supplemented with 10% v/v defibrinated sheep blood or in modified Stainer-Scholte medium (SS). Media were supplemented with gentamicin 50 μg/ml or streptomycin 200 μg/ml as needed. Bacterial strains and plasmids used in this work are listed in Table S1.

*Escherichia coli* (DH5α and S17-1) strains were cultured with lysogeny broth (LB) either in test tubes or on LB agar plates (1.5% agar). Gentamicin was added to the medium at 10 μg/ml final concentration when appropriate. Replicative plasmids were introduced to *E. coli* by electroporation using standard techniques. Non-and replicating plasmids were introduced into *B. bronchiseptica* by conjugation. The yeast strain InvSc1 (*Saccharomyces cerevisiae*; Invitrogen), was routinely cultured on YPD medium. To select plasmids carrying the URA3 gene, yeast was grown on YNB medium with a complete supplemental mixture minus uracil.

### Strain and plasmid construction

All oligonucleotide primers used in the study are listed in Table S2. Cloning was performed by *in vivo* recombination in yeast, as described by Shanks (21). Briefly, a plasmid derived from the pMQ30 allelic replacement vector was utilized to create knockout strains. Two homologous DNA regions flanking the target genomic area were amplified by PCR, designated F1 and F2, using primers with over 30 additional bases to aid in recombination with adjacent fragments. These fragments were inserted into the pMQ30 plasmid using yeast cloning methods, resulting in pMQ30F1F2. The plasmid was then recovered from yeast and electroporated into *E. coli* S17-1. All constructs were confirmed by PCR and DNA sequencing. Allelic replacement mutants were generated as previously described (22). In brief, the plasmid was introduced into wild type *B. bronchiseptica* via biparental conjugation. Simple recombinants were selected using streptomycin and gentamicin. For a detailed protocol of *B. bronchiseptica* conjugation, see reference (22). Clones resistant to both antibiotics were then grown in SS medium without antibiotics and plated on LB containing 11% sucrose to isolate clones that had lost the plasmid. These clones were checked for gentamicin sensitivity, and the deletion was confirmed by PCR using primers that flank the deleted region.

### Biofilm formation assays

*B. bronchiseptica* biofilm assays were performed as previously reported by our group (5). Cultures with an OD_650nm_ of 0.1 were prepared by resuspending colonies grown on SS (1.5% w/v agar) supplemented with 10% v/v defibrinated sheep blood into SS. One hundred microliters of these cultures were then pipetted into a 96-well U-bottom microtiter plate (polyvinylchloride, PVC) and incubated statically at 37°C for 24 hours. After incubation, planktonic bacteria were removed by washing, and attached bacteria were stained with a 0.1% w/v CV solution. The stain was then solubilized in 120 μl of a 33% v/v acetic acid solution. Biofilm formation was quantified by measuring Abs at 595 nm of 100 μl of the solubilized stain solution. Nicotinic acid was added at specified concentrations when used, and bovine serum albumin (BSA) and CaCl_2_ were added where indicated, at final concentrations of 2 mg/ml and 2.0 mM, respectively. The experiments were repeated at least three times, each with four technical replicates.

### Measurements of c-di-GMP levels

C-di-GMP levels were analyzed via LC-MS as previously described (5). Briefly, four replicates of each strain and condition were harvested and resuspended in 250 μl of extraction buffer (methanol-acetonitrile-water [40:40:20] plus 0.1 N formic acid at −20°C) and incubated at −20°C for 30 min. The cell debris was pelleted for 5 min at 4°C, and the supernatant containing the nucleotide extract was immediately adjusted to a pH of ∼7.5 with 15% (NH_4_)_2_HCO_3_ and stored on dry ice prior to analysis. The resultant extract was analyzed via LC-MS using the LC-20AD high-performance LC system (Shimadzu, Columbia, MD) coupled to a Finnigan TSQ Quantum Discovery MAX triple-quadrupole mass spectrometer (Thermo Electron Corp., San Jose, CA).

### Western blots

*B. bronchiseptica* strains were grown for 48 hours on BGA plates and then transferred to SS liquid culture for overnight growth. Samples were collected, and the OD_650nm_ was standardized. After centrifugation at 13,500 rpm for 3 minutes, the supernatants intended for HA tag detection were filtered through a low-binding protein 0.22 μm filter, transferred to new tubes, and concentrated using an Amicon Ultra centrifugal filter with a molecular weight cutoff (MWCO) of 10.0 kDa. Bacterial pellets were resuspended in PBS to achieve an OD_650nm_ of 10 for BrtA-HA detection.

The quantity of protein in the supernatants was measured with BCA Pierce Kit. All samples were boiled for 10 min and centrifuged before loading. Proteins were separated by SDS-PAGE 8% and transferred to a nitrocellulose membrane. BrtA-HA was detected with anti-HA diluted 1:2,000 in 3% w/v BSA in TBS overnight at 4 °C, followed by incubation with anti-mouse IgG conjugated to horseradish peroxidase (HRP) (1:15,000) (Invitrogen) in TBS-0,01% Tween containing 3% BSA at room temperature for 2.5 h. Detection was performed using Clarity™ Western ECL Substrate (Bio Rad #1705060) according to the manufacturer’s instructions.

### Dot Blot

*B. bronchiseptica* strains were grown as described above for western blot analysis. An aliquot of 1.5 ml was centrifuged at 16,000 x g for 3 min and the pellets were washed two times with 1 ml of buffer (Tris 20 mM, MgCl2 10 mM, pH = 8,0) and finally resuspended in the same buffer to obtain a bacterial suspension with an OD of 1. The bacterial suspension were alternatively diluted 2-fold three times (½, ¼, ⅛) and 10 μl of these samples were pipetted onto a nitrocellulose membrane, or pippetted directly in triplicate. Once dried, the membrane was incubated for 1 h with blocking buffer (5% w/v non fat milk in PBS 0,01% Tween 20). BrtA and FHA were detected using a mouse monoclonal anti-HA or anti-FHA antibody (1:5,000 in blocking buffer) as the primary antibody and a goat anti mouse or anti rabbit monoclonal antibody respectively conjugated to HRP as the secondary antibody. Non blocked samples were stained with Ponceau red to confirmed equal protein concentration of samples.

### Scanning electron microscopy

Biofilm assays for scanning electron microscopy were conducted as previously described by our group (5). *B. bronchiseptica* was cultured in SS medium on glass coverslips, which were vertically submerged in plastic tubes for 24 hours. The coverslips were then subjected to a CO_2_-critical-point procedure using an EmiTech K850 and sputter-coated with gold. Samples were examined with a scanning electron microscope (FEI Quanta 200), and images were processed using the Image Soft Imaging System ADDA II. At least two independent samples were analyzed per strain, with a majority of each sample scanned and representative images selected for processing.

### Statistical analysis

Each experiment included at least two biological replicates (as indicated in each experiment). Data were analyzed for statistical significance using a one-way ANOVA followed by Tukey’s multiple-comparison test to assess differences among groups. The significance level is specified in each figure legend.

## Data availability

The GenBank accession number for the *B. bronchiseptica* RB50 genome is NC_002927.3. The gene identification number and locus tag, respectively, for *bdcB* are 2661408 and BB_RS19575, *pdeA* are 2660213 and BB_RS13380, *pdeC* are 2661280 and BB_RS15655 and *pdeB* are 2661612 and BB_RS15715.

## Acknowledgments

This work was supported by grants from Argentina; PICT-2019-00680 and PID-UNLP-X1008 to J.F. and F.S. SLM is a fellow from ANPCyT (PICT-2019-00680). F.S. and J.F. are members of the Research Career of CONICET. All authors contributed to the discussion and provided comments on the manuscript. All authors approved the final version of the manuscript.

